# Surrogate R-spondins for tissue-specific potentiation of Wnt signaling

**DOI:** 10.1101/487223

**Authors:** Vincent C. Luca, Yi Miao, Xingnan Li, Michael J. Hollander, Calvin J. Kuo, K. Christopher Garcia

## Abstract

Secreted R-spondin1-4 proteins (RSPO1-4) orchestrate stem cell renewal and tissue homeostasis by potentiating Wnt/β-catenin signaling. RSPOs induce the turnover of negative Wnt regulators RNF43 and ZNRF3 through a process that requires RSPO interactions with either Leucine-rich repeat containing G-protein coupled receptors (LGRs) or heparin sulfate proteoglycans (HSPGs). Here, we describe the engineering of ‘surrogate RSPOs’ that function independently of LGRs and HSPGs to enhance Wnt signaling on cell types expressing a target surface marker. These bispecific proteins were generated by fusing an RNF43- or ZNRF3-specific single chain antibody variable fragment (scFv) to the immune cytokine IL-2. Surrogate RSPOs mimic the function of natural RSPOs by crosslinking the extracellular domain (ECD) of RNF43 or ZNRF3 to the ECD of the IL-2 receptor CD25, which sequesters the complex and results in highly selective amplification of Wnt signaling on CD25+ cells. Furthermore, surrogate RSPOs were able substitute for wild type RSPO in a colon organoid growth assay when intestinal stem cells were transduced to express CD25. Our results provide proof-of-concept for a technology that may be adapted for use on a broad range of cell- or tissue-types and will open new avenues for the development of Wnt-based therapeutics for regenerative medicine.

## INTRODUCTION

The Wnt/β-catenin pathway controls cell fate determination and tissue homeostasis in all metazoans^1^. Pleiotropic Wnt signaling exhibits differential effects on a wide array of cell types and is therefore tightly regulated by several host-encoded enhancers and inhibitors. R-spondin proteins (RSPO1-4 in mammals) potentiate Wnt signaling by antagonizing negative regulators of the Wnt receptor Frizzled^2^. The effect of RSPO is remarkably potent, and co-administration of Wnts and RSPOs can result in signaling outputs that are several hundred-fold greater than those of Wnt alone^3^.

RSPO-mediated enhancement occurs via an indirect mechanism that greatly increases expression levels of the Wnt receptors Frizzled (Fzd) and LRP5 or LRP6 on the cell surface. In the absence of RSPO, the transmembrane E3 ligases RNF43 and ZNRF3^4^ ubiquitinate the intracellular regions of Fzd, which results in the internalization and degradation of both Fzd and associated LRP5/6 proteins (Fig 1A). RSPO proteins are comprised of two Furin-like domains (Fu1, Fu2) followed by a thrombospondin domain (TSP) and basic region (BR). The Fu1 domain of RSPO binds to the extracellular domains (ECDs) of RNF43 or ZNRF3 while the Fu2 domain binds to the ECD of a co-receptor, the Leucine-rich repeat-containing G-protein coupled receptor 4, 5 or 6 (LGR4-6)^5-8^. This crosslinking event induces endocytosis of the RSPO-LGR-E3 ligase ternary complex, which sequesters RNF43 or ZNRF3 from Fzd and, in turn, potentiates Wnt signaling (Fig 1B). It was recently determined that RSPO2 and RSPO3 can function independently of LGRs by binding to heparan sulfate proteoglycans (HSPGs) on the cell surface via their TSP and BR domains^9^. These data suggest RSPO are modular molecules consisting of an RNF43/ZNRF3 binding domain that is necessary for the potentiation of Wnt signaling and an LGR/HSPG-binding domains that facilitates internalization of the complex.

**Figure 1.**
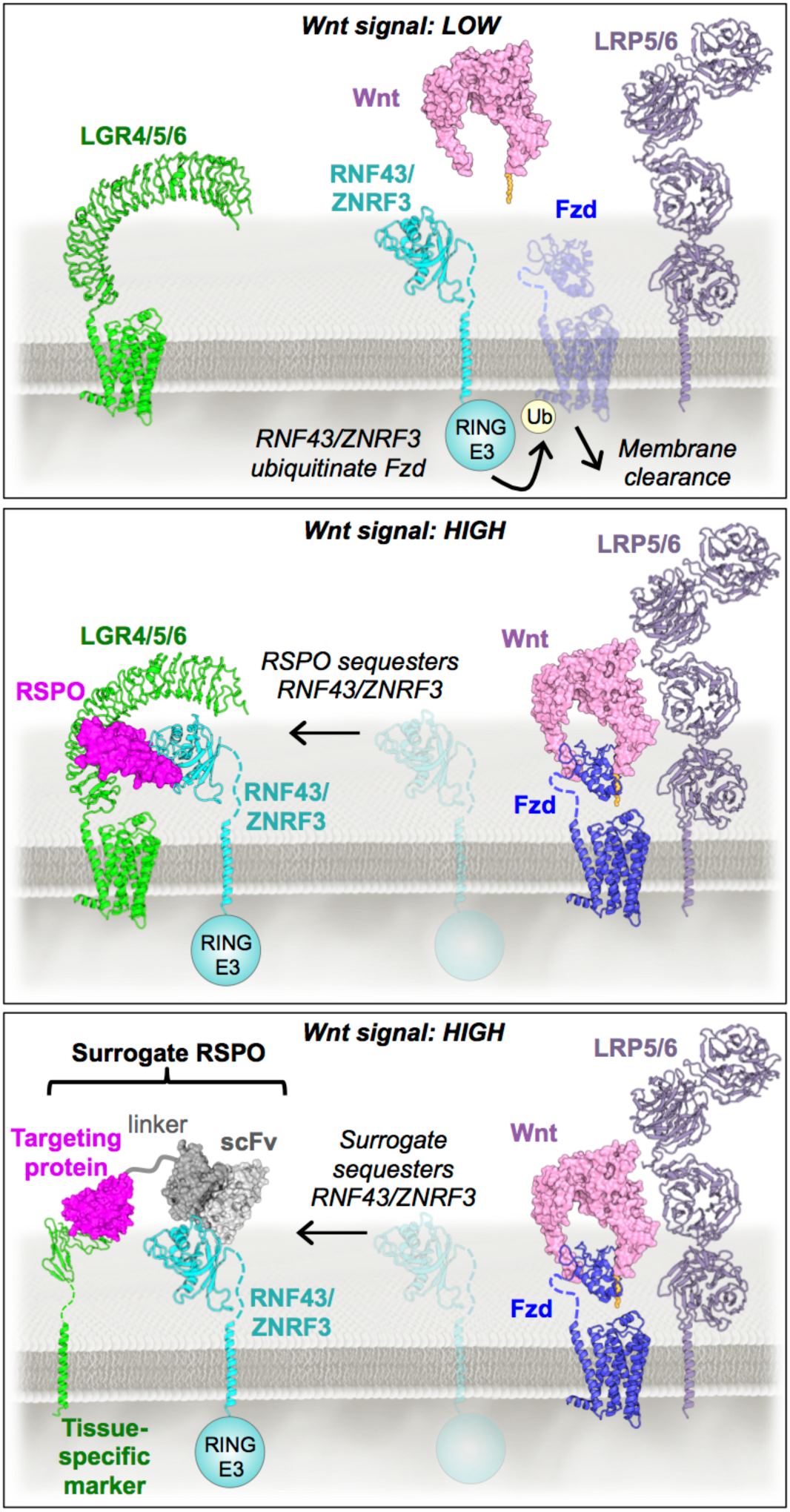
The R-spondin signaling mechanism. In the absence of RSPO, ZNRF3 drives membrane clearance of Fzd receptors to negatively regulate Wnt signaling. RSPO-mediated crosslinking of the ECDs of RNF43 or ZNRF3 and LGR4, LGR5, or LGR6 sequesters RNF43/ZNRF3 to restore Fzd surface levels, thereby potentiating Wnt activity. Surrogate RSPOs mimic the function of wild type RSPOs by cross-linking RNF43 or ZNRF3 to tissue-specific markers known to undergo endocytosis upon ligand binding. PDB files 5UK5^7^, 4F0A^25^, 3S8Z, and 3S94^26^ were used to generate the images above.

Biomedical applications of recombinant RSPOs have been widely explored in the field of regenerative medicine, especially in the context of the small intestine where RSPOs function to promote the renewal of stem cells at the base of the crypt^10,11^. For example, recombinant RSPO1 protects animals from both experimentally induced colitis and from lethal doses of chemoradiation by stimulating intestinal regeneration^12^. Furthermore, the addition of exogenous RSPO is critical for the proliferation of intestinal organoids^10^, an *in vitro* model system for intestinal epithelium studies. Interestingly, it has been recently reported that Wnts and RSPOs play non-equivalent roles in intestinal stem cell (ISC) self-renewal through a feedback loop, in which Wnts drive the expression of LGR5 and RSPOs induce ISC expansion^13^, raising the possibility that targeting the pathways individually or in concert may be associated with unique therapeutic outcomes.

Potential obstacles for the clinical use of Wnt agonists include toxicity, off-target effects, and diminished potency caused by RNF43/ZNRF3-mediated antagonism. Wnts are powerful morphogens and hyperactivation of the Wnt pathway has been implicated in the development of several human cancers^14^. RSPOs are promising candidates to selectively boost innate Wnt activity while circumventing off-target effects associated with global Wnt agonism. However, our ability to target RSPOs to specific tissue or cell types is restricted by the expression profiles of RSPO receptors, since HSPGs are ubiquitously expressed and LGR expression is restricted to subsets of tissue stem cells. To develop precisely targeted RSPO mimetics, we engineered synthetic protein ligands that phenocopy RSPO-mediated Wnt signal enhancement via an LGR- and HSPG-independent mechanism. These “surrogate RSPOs” function by cross-linking the RNF43 or ZNRF3 ECD to an immune cell surface marker (CD25) that is known to undergo internalization upon ligand binding^15^, thereby mimicking RSPO-mediated sequestration of RNF43/ZNRF3 (Fig. 1). Surrogate RSPOs selectively boost Wnt signaling in CD25+ reporter cells and human colon organoids, and their modular design allows them to be adapted to target virtually any cell type, a feature that has major implications for the development of Wnt-based therapeutics.

## RESULTS

### Isolation of RNF43- and ZNRF3-specific scFvs using yeast surface display

Our surrogate RSPO design concept involves the fusion of two components: (i) a RNF43- or ZNRF3-binding protein and (ii) a tissue-specific targeting protein that binds to a surface receptor. Administration of the surrogate RSPO protein will cross-link ECD of RNF43 or ZNRF3 to the target and mimic the function of natural RSPOs by sequestering or internalizing the complex. We isolated the RNF43- or ZNRF3-binding protein components for the surrogate RSPOs by selecting single-chain variable fragments (scFvs) from a published yeast display library of ∼1×10^9^ sequences derived from human B-cells (Fig. 2A)^16^. The scFvs were obtained by performing two parallel sets of selections against either the RNF43 ECD or the ZNRF3 ECD using magnetic activated cell sorting (MACS) and fluorescence activated cell sorting (FACS). After several rounds of selection, we isolated a panel of different RNF43- or ZNRF3-specific scFvs (Fig. S1), and the clones “R5” and “Z6” were chosen for incorporation into surrogate RSPOs based on their ability to robustly bind to RNF43 and ZNRF3, respectively, in a fluorescence-based assay (Fig. 2B).

**Figure 2.**
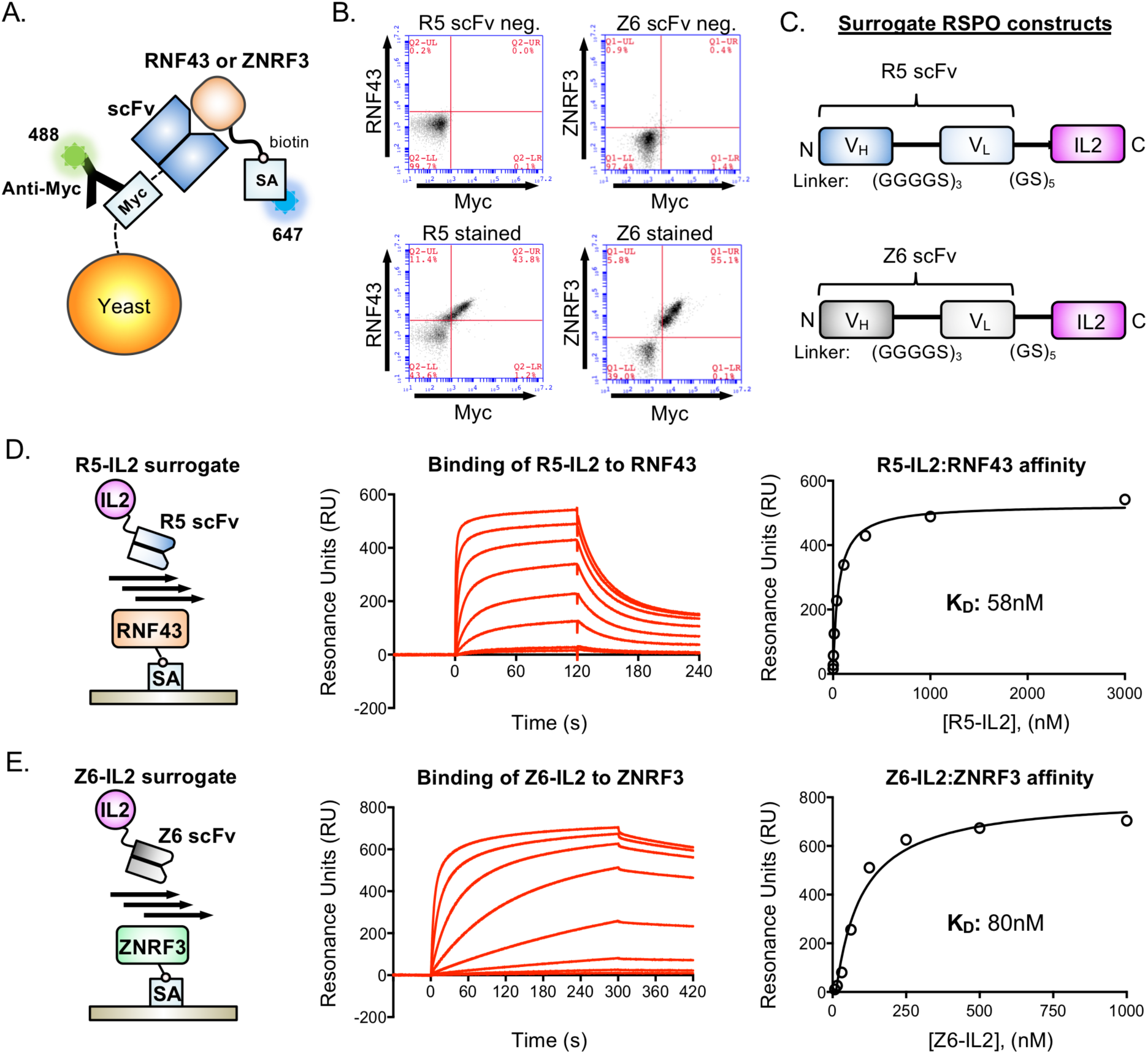
Characterization of RNF43- and ZNRF3-specific scFvs used for generation of surrogate RSPOs. (a) Cartoon schematic of yeast-displayed scFv binding to biotinylated RNF43 or ZNRF3 ECDs. (b) Flow cytometry dot plots depicting the binding of R5 and Z6 scFvs to RNF43 and ZNRF3, respectively. R5- or Z6-expressing yeast were stained with 1 µM concentrations of biotinylated RNF43 or ZNRF3 ECDs, respectively, and binding was detected using fluorescently labeled streptavidin. Surface expression of R5 and Z6 was detected with an antibody to the c-Myc epitope. Negative controls are unstained R5- or Z6-expressing yeast. (c) Construct design for R5-IL2 and Z6-IL2 surrogate RSPOs. (d) & (e). SPR was used to measure the binding of R5-IL2 or Z6-IL2 to RNF43 and ZNRF3, respectively. Cartoons (left) depict the orientation of streptavidin (SA)-coupled RNF43 or ZNRF3 ECDs and the R5-IL2 or Z6-IL2 analytes on the sensor chip. Curves (center) indicate a series of injections of R5-IL2 (3-fold dilutions, maximum concentration 3 µM) or Z6-IL2 (2-fold dilutions, maximum concentration 1 µM). Dissociation constants were obtained by plotting maximum RU values and fitting to a 1:1 binding model (right).

### Design, expression and purification of surrogate RSPOs

Membrane clearance of RSPO signaling complexes has been reported to require the internalization of the receptors LGR4-6 ^17^. We therefore selected the immune cytokine IL-2 to act as a proxy for RSPO-LGR binding in our surrogate RSPO constructs based on its ability to induce endocytosis of its cognate receptor, CD25 ^15^. We generated two surrogate RSPO constructs, one targeted to RNF43 and one targeted to ZNRF3, by fusing R5 to IL-2 (R5-IL2) and Z6 to IL-2 (Z6-IL2). Single-chain constructs were designed such that R5 or Z6 is connected to the N-terminus of IL-2 via a flexible 5x(Gly-Ser) linker (Fig. 2C), which allows R5/Z6 and IL-2 to independently bind their respective targets. R5-IL2 and Z6-IL2 were expressed in insect cells using Baculovirus and both proteins eluted from a gel filtration column as monodisperse peaks, which indicates favorable biochemical behavior (Fig. S2).

### Biophysical characterization of surrogate RSPO-receptor interactions

We used surface plasmon resonance (SPR) to measure the binding affinity of R5-IL2 and Z6-IL2 for RNF43 and ZNRF3, respectively. We determined that R5-IL2 bound to RNF43 with a dissociation constant (K_d_) of 58 nM, and that Z6-IL2 bound to ZNRF3 with a K_d_ of 80 nM (Fig. 2D and 2E). R5-IL2 did not cross-react with ZNRF3, and Z6-IL2 did not cross-react with RNF43 (Fig. S3). We also used SPR to determine that R5-IL2 bound to CD25 with a Kd of 8.4 nM and that Z6-IL2 bound to CD25 5.7 nM, both of which values are similar to the reported K_d_ values of the wild type CD25-IL2 interaction (Fig. S4)^18,19^. Collectively, our SPR experiments revealed that (i) R5 and Z6 bind to their respective targets with similar affinities, (ii) that the affinities of surrogate RSPOs fall within the range of those reported between wild type RSPOs and RNF43/ZNRF3 (5 nM to 10,000 nM) 7,8, and (iii) that CD25 binding is retained in the surrogate RSPO format.

### Surrogate RSPO-mediated enhancement of Wnt signaling

We performed a luciferase reporter assay to test the selectivity of R5-IL2 and Z6-IL2 in potentiating Wnt signaling. We compared the activity of surrogate RSPOs on HEK 293 Super Top Flash (STF) Wnt reporter cells, or on HEK 293 STF cells that had been retrovirally transduced with CD25. In the CD25-expressing cells, R5-IL2 increased WNT3a reporter activity up to 9.5-fold, and Z6-IL2 increased WNT3a activity up to 41-fold (Fig. 3A). The greater activity of Z6-IL2 relative to R5-IL2 is consistent with a previous report that ZNRF3 is the dominantly expressed homolog (relative to RNF43) in HEK 293 cells^2^. Given that natural RSPOs are cross-reactive, and that sequestration of both RNF43 and ZNRF3 may be required for the potent activity of RSPOs, we posited that R5-IL2 and Z6-IL2 would have increased potency when co-administered. We therefore incubated the cells with a 1:1 mixture of R5-IL2 and Z6-IL2, and found that this combination increased WNT3a activity by 148-fold (Fig. 3A). However, in the untransduced cells, addition of R5-IL2, Z6-IL2, and R5-IL2+Z6-IL2 led to increases in WNT3a activity of only 2.7, 3.9 and 5.8-fold, respectively (Fig. 3B), indicating that surrogate RSPO activity is highly selective for cells expressing CD25.

**Figure 3.**
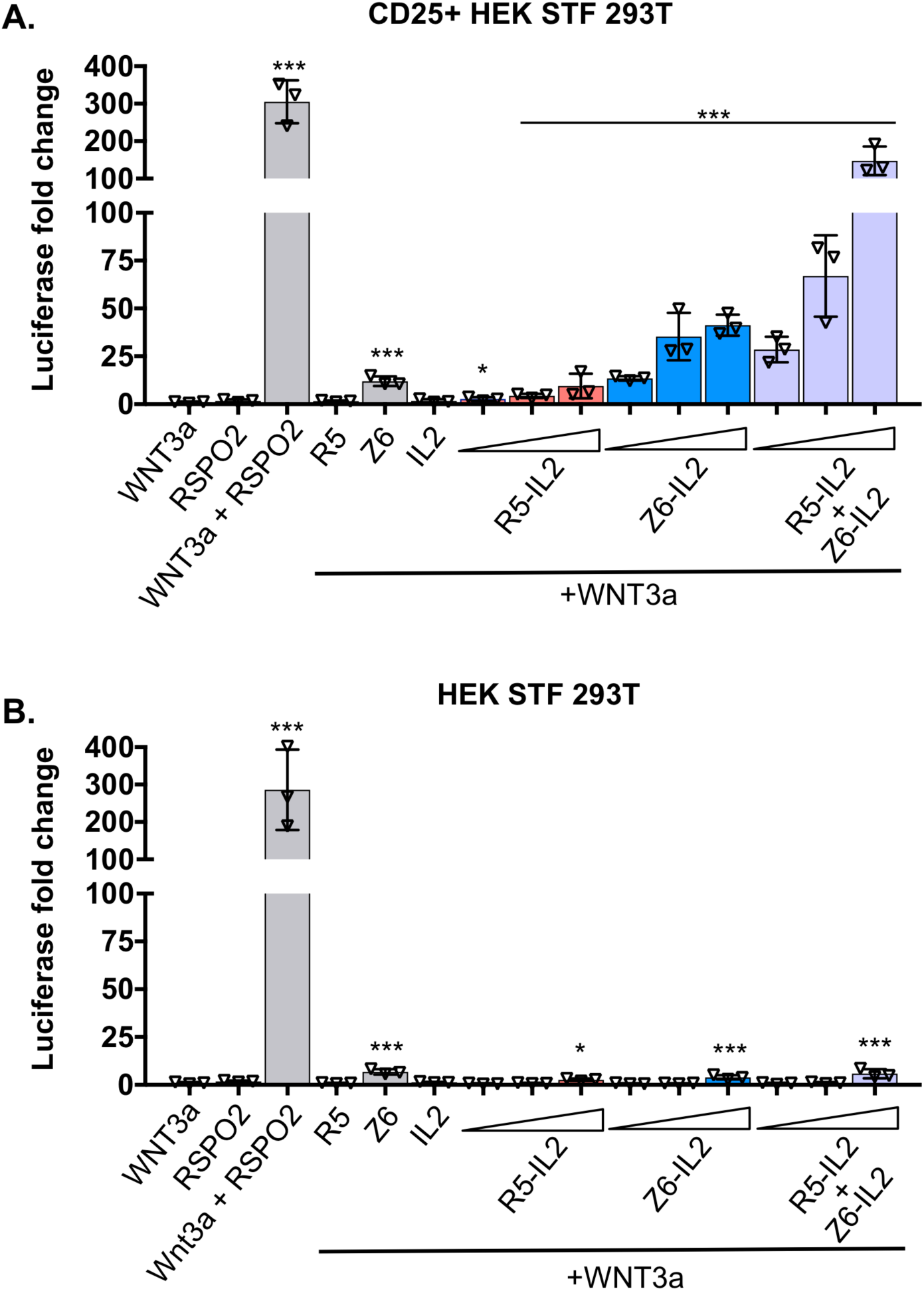
Surrogate RSPOs potentiate Wnt signaling. (A) Representative Luciferase reporter assay measuring surrogate RSPO potentiation of Wnt activity in CD25-expressing cells. CD25-expressing HEK STF 293T cells were treated 20% WNT3a conditioned media and various recombinant proteins (RSPO2 Furin domains 1 and 2 (25 nM), R5 (500 nM), Z6 (500 nM), IL-2 (500 nM), R5-IL2 (5 nM, 50 nM, 500 nM), Z6-IL2 (5nM, 50 nM, 500 nM), or a 1:1 mixture of R5-IL2 and Z6-IL2 (5 nM each, 50 nM each, 500 nM each). The Luciferase signal fold change is normalized to WNT3a alone. Error bars represent standard deviation of n=3 technical replicates. Statistical significance of WNT3a alone versus each condition was determined by using one-way ANOVA on log-transformed data. * P<0.05, ** P<0.01, *** P<0.001. (B) A reporter assay was performed under the same conditions as in (A) using uninfected (CD25 negative) HEK STF 293T cells. * P<0.05, ** P<0.01, *** P<0.001.

In both CD25 positive and CD25 negative cells, we found that a mixture of conditioned WNT3a media and RSPO2 (Furin domains 1 and 2) elicited a response that was 286- to 304-fold greater than that of WNT3a alone (Fig. 3A, 3B), which is similar to the previously observed levels of RSPO-mediated enhancement ^3^. As negative controls, we treated cells with WNT3a conditioned media supplemented with either R5, Z6, or IL-2. The R5 scFv and IL2 cytokine did not substantially potentiate WNT3a signaling (Fig. 3B), however, the Z6 scFv exhibited mild Wnt-potentiating activity that resulted in 10-fold or 12-fold increases in signaling in the HEK293 STF and CD25-expressing cells, respectively (Fig. 3A, 3B).

### Surrogate RSPO-mediated stimulation of intestinal organoid growth

The growth of primary human colon organoids is presumably driven by LGR5+ stem cells and requires the addition of exogenous RSPOs along with Wnt, EGF and Noggin^8,9^. To determine whether R5-IL2 or Z6-IL2 can substitute for RSPOs in human colon organoid culture, we performed an organoid growth assay using organoids that had been retrovirally transduced to express CD25. In this assay, CD25+ organoid cells were cultured in a basal medium of WNT3a, EGF and Noggin, supplemented with one of the following recombinant proteins: RSPO2, IL-2, R5 scFv, Z6 scFv, R5-IL2, Z6-IL2, or a 1:1 mixture of R5-IL2 and Z6-IL2, Subsequently, organoid growth was monitored by resazurin fluorescence in a cell viability assay. Although not as potent as RSPO2 (73-fold), addition of R5-IL2, Z6-IL2, or the mixture of R5-IL2 and Z6-IL2 each led to significant increases in organoid growth relative to the negative control protein, IL-2 (R5-IL2: 7.3-fold, Z6-IL2: 9.4-fold, mixture: 11-fold) (Fig. 4). However, addition of R5 scFv or Z6 scFv did not result in a significant increase. Importantly, neither R5-IL2 nor Z6-IL2 stimulated the growth of wild type organoids (Fig. 4), indicating that surrogate RSPOs specifically affect cells expressing CD25.

**Figure 4.**
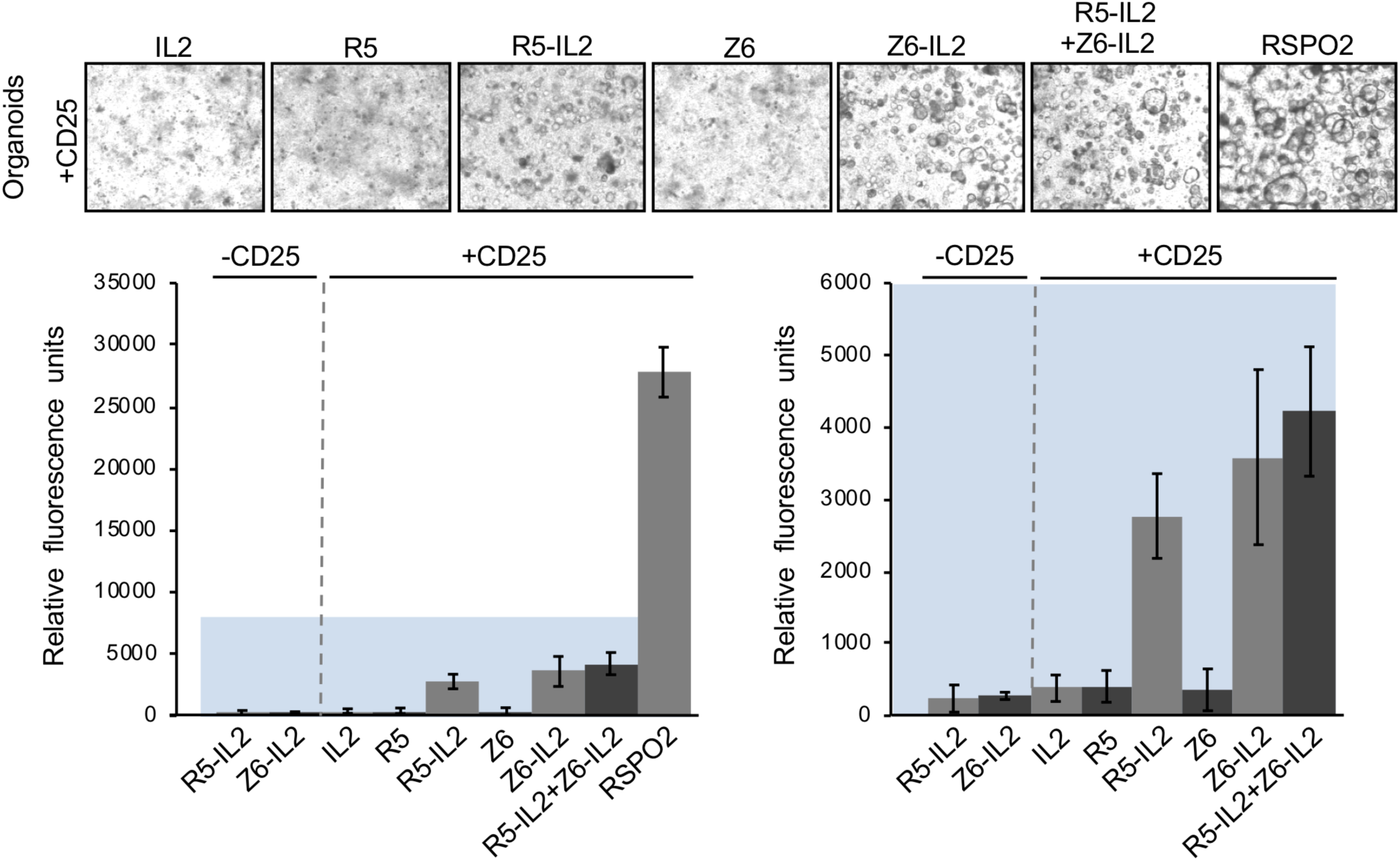
Surrogate RSPOs stimulate intestinal organoid growth. Surrogate RSPOs were tested for their ability to stimulate the growth of LGR5+ human colon organoids that had been retrovirally transduced to express CD25. Basal media containing WNT3a, EGF and Noggin, but lacking RSPO2 was supplemented with 500nM concentrations of the indicated proteins, or with 25nM RSPO2. Microscopy was used to observe the appearance of spherical organoids in each condition (top) and growth was quantitated by resazurin fluorescence in a cell viability assay (bar graphs, bottom). The bar graph on the right is a zoomed panel of the colored region from the graph on the left. Error bars represent standard error of n=12.

## DISCUSSION

The successful development of surrogate RSPOs indicates that a non-natural receptor, CD25, may be co-opted to selective amplification of Wnt signaling. The bispecific R5-IL2 and Z6-IL2 proteins do not include any components derived from natural RSPOs and do not recognize LGR receptors or HSPGs. The Wnt-potentiating activity of surrogate RSPOs is potentially driven by IL-2-mediated endocytosis of CD25, which would support the present model that RSPOs function by inducing membrane clearance of RNF43 and ZNRF3. However, it is also possible that surrogate RSPOs sequester RNF43 and ZNRF3 via an unknown mechanism that parallels the recently discovered HSPG-driven activity of RSPO2 and RSPO3. Regardless, the development of R5-IL2 and Z6-IL2 will open new avenues for the engineering surrogate RSPOs that act upon a broad spectrum of cell surface receptors. For example, tyrosine kinase receptors, G-protein coupled receptors (GPCRs), pattern recognition receptors and several other classes of receptors are internalized upon ligand-binding and may serve as alternatives for the targeting arms of surrogate RSPOs.

The selective amplification of Wnt signaling on CD25+ cells by R5-IL2 and Z6-IL2 has the potential to influence immune cell behavior for biomedical applications. CD25 is expressed predominantly on activated T cells, regulatory T cells (T regs), and natural killer cells (NK cells)^20^ and Wnt activation is known to modulate the behavior of several T cell subtypes. For example, Wnt signaling attenuates the suppressive activity of T regs by disrupting Foxp3 transcription activity^21^, and autocrine inhibition of Wnt signaling in NK cells mediates tumor immune evasion by downregulating the expression of activating ligands ^22^. Administration of CD25-targeted surrogate RSPOs alone or in combination with Wnt agonists may therefore augment the effectiveness of T cell- or NK-cell based cancer immunotherapies.

We designed R5-IL2 and Z6-IL2 such that they bind to RNF43 and ZNRF3, respectively, and neither surrogate RSPO is cross-reactive (Figs. 2D, 2E, S3). In our 293-based Wnt reporter assay, the highest levels of Wnt signal enhancement were achieved by co-administering R5-IL2 and Z6-IL2 (Fig. 3), suggesting that maximal activity requires the antagonism of both RNF43 and ZNRF3. Natural RSPOs, which are cross-reactive with both RNF43 and ZNRF3, were also more potent stimulators of colon organoid growth when compared to R5-IL2 and Z6-IL2 (Fig. 4). Based on these findings, we anticipate that second-generation surrogate RSPOs may be optimized through the incorporation of cross-reactive RNF43/ZNRF3 binding modules, such as the RSPO Fu1 domain^8^, in place of the scFv component. Additional optimization strategies may involve the use of scFvs that recognize different epitopes on the RNF43 or ZNRF3 ECDs, or the use of scFvs that bind to RNF43 or ZNRF3 with higher affinity. On the other hand, the ability of R5-IL2 and Z6-IL2 to target a given E3 ligase (RNF43 versus ZNRF3) confers an additional degree of selectivity that is not attainable using natural RSPOs, and this property may be advantageous for targeting cell types that express only one of the two E3 ligases. Collectively, our data provide proof-of-concept for surrogate RSPOs whose modular design will allow for the targeted enhancement of Wnt signaling on cells expressing RNF43, ZNRF3 or various other tissue-specific markers.

## METHODS

### Protein expression and purification

The ECDs of human RNF43 (amino acids 24-197) and ZNRF3 ECD (amino acids 56-219) were cloned into the pAcGp67A vector with a C-terminal biotin-acceptor peptide tag (GLNDIFEAQKIEW) followed by a hexahistidine tag. Unless otherwise specified, RNF43/ZNRF3 in the methods section will refer to their ECD only. Selected RNF43 and ZNRF3 high-affinity human antibody scFv fragments were cloned into pAcGp67A vector with a C-terminal hexahistidine tag. These human antibody scFv fragments were also cloned into pAcGp67A vector in frame with a (GS)_5_ linker and human interleukin-2 (amino acids 21-153) followed by a hexahistadine tag. All proteins were expressed in Hi-Five cells (Invitrogen) from *Trichoplusia ni* using Baculovirus and the cultures were harvested 60 hours after infection. Proteins were purified by nickel affinity chromatography followed by size exclusion chromatography in HBS buffer (10 mM HEPES pH 7.2, 150 mM sodium chloride). RNF43 and ZNRF3 were site-specifically biotinylated by BirA ligase at the C-terminal biotin-acceptor peptide prior to size-exclusion chromatography. All proteins were either used when freshly purified or snap frozen in liquid nitrogen with 20% glycerol and then thawed for use as necessary.

### Selection of RNF43/ZNRF3-binding scFv fragments

The nonimmune yeast surface display library of human antibody scFv fragments was generously provided by the group of Prof. Dane Wittrup (MIT) ^16^. A sequential selection strategy involving a series of magnetic bead selections followed by flow-cytometric sorting was used as previously reported^23^. Round 1 of the selections was performed by mixing 250 µL magnetic streptavidin microbeads (Militenyi) with 400 nM biotinylated RNF43/ZNRF3. These microbeads were further mixed with 1 × 10^10^ yeast cells from the scFv library ^16^. Yeast were subsequently passed over a Magnetic-Activated cell sorting (MACS) LS separation column (Militenyi), the flow through was discarded and ZNRF3/RNF43 binders was isolated from the column elution. Round 2 was performed as Round 1 except 1 × 10^8^ yeast cells from the recovered Round 1 sample were used for the selection. For Round 3, yeast cells incubated with 200 nM biotinylated RNF43 or ZNRF3 ECD, washed and then stained with streptavidin labeled with 50 nM Alexafluor-647 (SA-647). The ZNRF3/RNF43 binders were enriched by MACS using anti-647 microbeads (Militenyi). In Round 4, yeast cells were pre-incubated with 5 nM biotinylated RNF43 or 10 nM biotinylated ZNRF3, washed, and then double-stained with SA-647 and Alexa Fluor 488-conjugated antibody to the c-Myc epitope (Myc-488, Cell Signaling). High-affinity RNF43/ZNRF3 binders were isolated by fluorescence-activated cell sorting (FACS).

Plasmids from the final round of selection were isolated using the Zymoprep Yeast Plasmid Miniprep Kit (Zymo Research) and sequenced. These plasmids were electroporated into *S. cerevisiae* EBY100 yeast and recovered in SDCAA selection media, followed by induction in SGCAA induction media^23^. The individual scFv-transformed yeast was incubated with increasing concentrations of biotinylated RNF43 or ZNRF3 proteins. These were stained with SA-647 and Myc-488 and fluorescence was monitored by flow cytometry. The data were analyzed by GraphPad Prism 7 and the highest affinity clones encoding full scFv sequences (R5, Z6) were selected for use in subsequent experiments.

### Binding of RNF43 and ZNRF3 to yeast surface displayed scFvs

Yeast cells expressing either the R5 or Z6 scFvs were stained with 1 µM recombinant biotinylated RNF43 or ZNR3 ECDs, respectively, in phosphate buffered saline (PBS) (Thermo Fisher Scientific) + 0.5% BSA + 2 mM EDTA at 4 °C for 2 hours with slow rotation. Cells were then washed, incubated with SA-647 and Myc-488 for 20 min, washed again, and then analyzed by flow cytometry.

### Surface plasmon resonance

All binding measurements were conducted using a BIAcore T100 instrument (GE Healthcare). Biotinylated RNF43, ZNRF3 and CD25 were coupled on a SA sensor chip (GE Healthcare) at low density. An unrelated biotinylated protein was captured at equivalent coupling density to the control flow cells. Increasing concentrations of R5-IL2 and Z6-IL2 were injected onto the chip in HBS-P (GE Healthcare) at 30µl/mL for 120 s or 300 s, respectively. Subsequently, the chip was regenerated with 0.5 M MgCl_2_ and 25% ethylene glycol in HBS-P buffer for 80 s after each injection. Resonance units were calculated by subtracting the resonance units observed in RNF43, ZNRF3, or CD25-containing flow cells from those of the control flow cell. Curves were fitted to a 1:1 binding model using the accompanying Biacore evaluation software (Biacore/GE Healthcare).

### Retrovirus production

Human full length CD25 was cloned into retroviral vector pMSCV-IRES-YFP (gift from Dr. Melissa McCracken, Stanford University). Retrovirus was packaged in HEK293T cells as described^24^. Briefly, HEK293T cells were plated the day before transfection. Cells were transfected with pMSCV-CD25-IRES-GFP and pCL-10A packaging vector using X-tremeGENE HP (Sigma-Aldrich). The supernatant was collected, filtered using sterile 0.45 um filter.

### Luciferase signaling assays

HEK293T cells were stably transduced with Firefly Luciferase Reporter under the control of a concatemer of seven LEF/TCF binding sites and Renilla Luciferase Reporter by lentivirus as previously reported from the Garcia laboratory^3^. These cells were further stably infected with human CD25 by retrovirus co-expressing YFP. Transduced HEK293T cells were FACS sorted by YFP expression. CD25 surface expression were further confirmed by staining the transduced HEK293T cells with anti-CD25 Brilliant Violet 605 antibody (BioLegend, Clone BC96, #302632). Fluorescence was monitored by flow cytometry and was compared to non-transduced HEK293T cells (data not shown). The assays were conducted using the Dual Luciferase Assay kit (Promega) as instructed. Briefly, cells were plated at a density of 10,000 cells per well in a 96-well plate and incubated for 24 hours prior to stimulation. Following the plating step, cells were stimulated with 20% WNT3a conditioned media (derived from WNT3a L cells, ATCC) supplemented with RSPO2 Furin1 and Furin 2 domains (25 nM), R5 (500 nM), Z6 (500 nM), IL-2 (500 nM), R5-IL2 (5, 50, or 500 nM), Z6-IL2 (5, 50, or 500nM), or a 1:1 mixture of R5-IL2 and Z6-IL2 (5 nM, 50 nM, or 500 nM of each protein). Cells were cultured in the presence of signaling proteins for another 24 hours before being washed and lysed as suggested by the instructions in the Dual Luciferase Assay Kit (Promega) manual. Luminance signals were recorded using SpectraMax Paradigm and analyzed by GraphPad Prism 7. Statistical significance of WNT3a alone versus each condition was determined by using one-way ANOVA on log-transformed data.

### Organoid growth assays

Fresh human colonic samples were collected at Stanford Hospital with informed consent and grown as primary organoids in Matrigel domes as described^3^. The culture was passaged several times in WENR media (WNT3a, R-spondin, EGF, Noggin) to confirm robust outgrowth of organoids. To evaluate surrogate activity, the organoids were transduced with retrovirus expressing CD25 and FACS sorted for YFP+ (YFP co-expression with CD25) to enrich CD25+ population. CD25+ cells were then subcultured in basal media containing WNT3a, EGF and Noggin but lacking RSPO2, and then supplemented with candidate surrogate R-spondins or RSPO2 as in Fig. 4. Organoid morphology was observed and phase contrast pictures were taken using Nikon TS100. To quantify cell growth, organoids were dissociated to single cells and plated at 10,000 cells per well in 96-well format. A fluorescence-based cell viability assay was performed 3 days after plating using AlamarBlue cell viability reagent (Thermo Fisher). Fluorescent signals were normalized to background signals from wells plated with Matrigel only.

### Use of tissues from human participants

Primary human tissues were obtained through the Stanford Tissue Bank from patients undergoing surgical resection at Stanford University Medical Center (SUMC). All experiments utilizing human material were approved by the SUMC Institutional Review Board and performed under protocols #28908, #26213 and #17425. Written informed consent for research was obtained from adult donors prior to tissue acquisition.

## Acknowledgements

We thank Lora Picton for edits and helpful comments. We thank Richard Reich from the Moffitt Cancer Center Biostatistics Core for advice with statistical analysis. This work was supported by a gift from the Mathers Fund, and the Ludwig Cancer Foundation (K.C.G.); the Howard Hughes Medical Institute (K.C.G.); NIH R01-DK115728(K.C.G. and C.J.K.) and NIH DK085527 (C.J.K).

## Author contributions

V.C.L. and Y.M. generated surrogate RSPO constructs, purified the proteins, and conducted luciferase signaling assays. V.C.L. performed SPR binding assays. Y.M. performed yeast display scFv selections and generated stable cell lines. X.L. performed human colon organoid growth assays. M.J.H. provided IL-2 and CD25 proteins and CD25 retrovirus. V.C.L., Y.M., X.L., C.J.K. and K.C.G. interpreted the results. V.C.L., Y.M. and K.C.G. wrote the manuscript. V.C.L. and K.C.G. designed the study.

## Data Availability Statement

The datasets generated during and/or analyzed during the current study are available from the corresponding author on reasonable request.

