## Supplementary Information for "Surrogate R-spondins for tissue-specific potentiation of Wnt signaling"

**Materials included:**

Figure S1

Figure S2

Figure S3

Figure S4

S-1

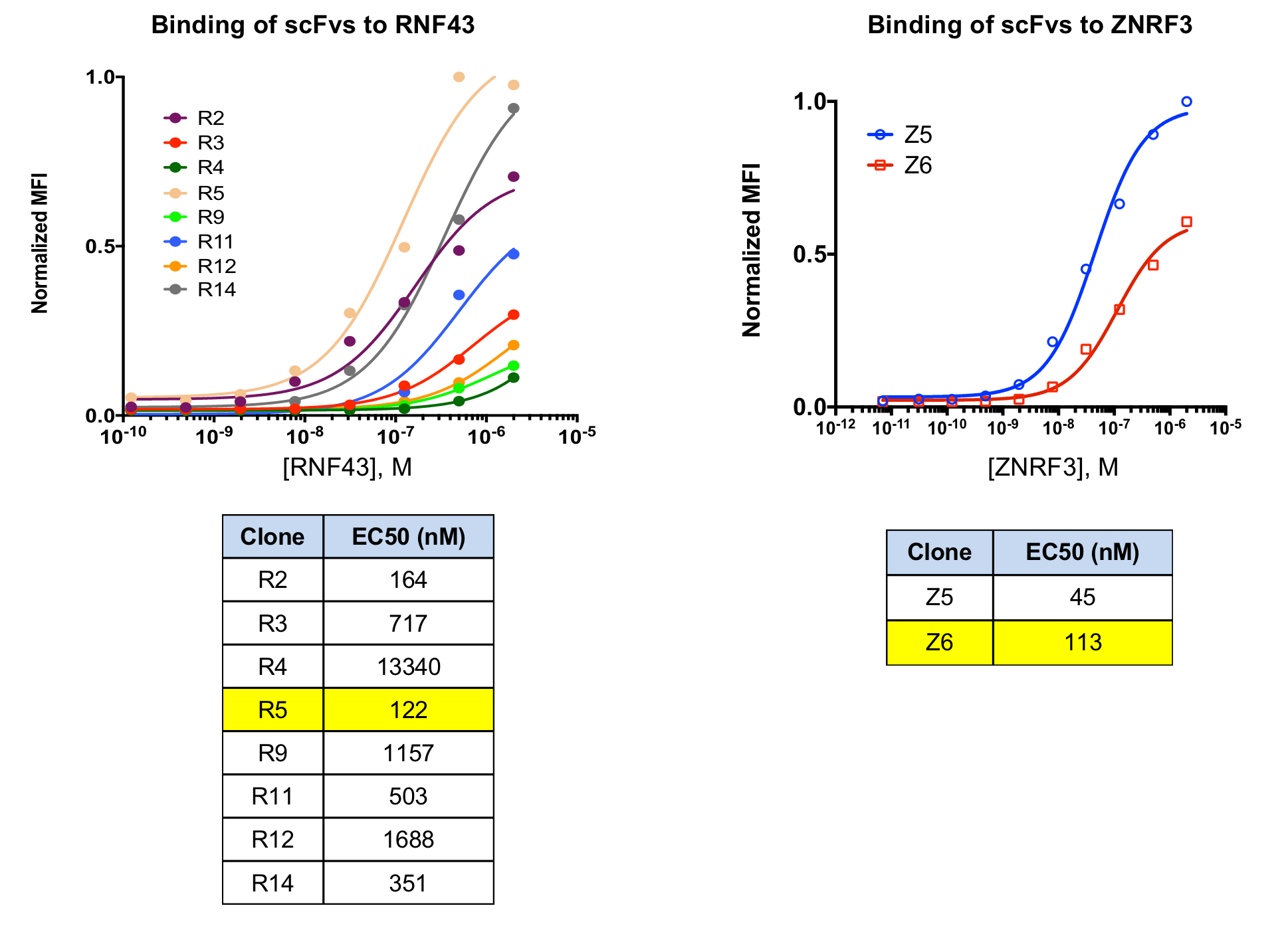

**Figure S1. Dose-response titrations of yeast displayed RNF43- and ZNRF3-specific scFvs.** Individual clones from the yeast scFv library were titrated with increasing concentrations of RNF43 (for RNF43-selected yeast) or ZNRF3 ECDs (for ZNRF3-selected yeast) and curves were fitted to determine EC50 values (indicated in charts below). The highlighted clones were selected for incorporation into surrogate RSPOs. The Z5 clone was excluded from future studies because it encoded for a truncated scFv sequence.

S-2

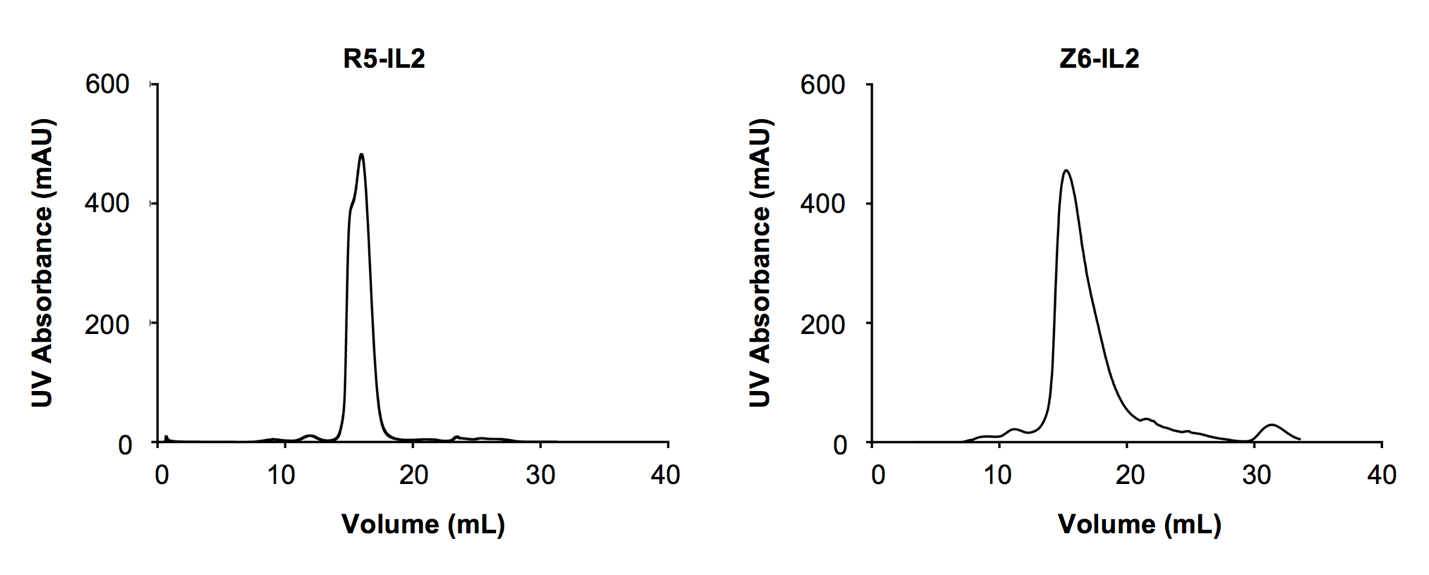

**Figure S2. Gel filtration chromatography of surrogate RSPOs.** R5-IL2 (left) and Z6-IL2 (right) were injected onto Sephadex 200 gel filtration columns and UV_280_ absorbance was plotted versus elution volume. Both proteins eluted predominately as monodisperse peaks, which is indicative of favorable biochemical behavior.

S-3

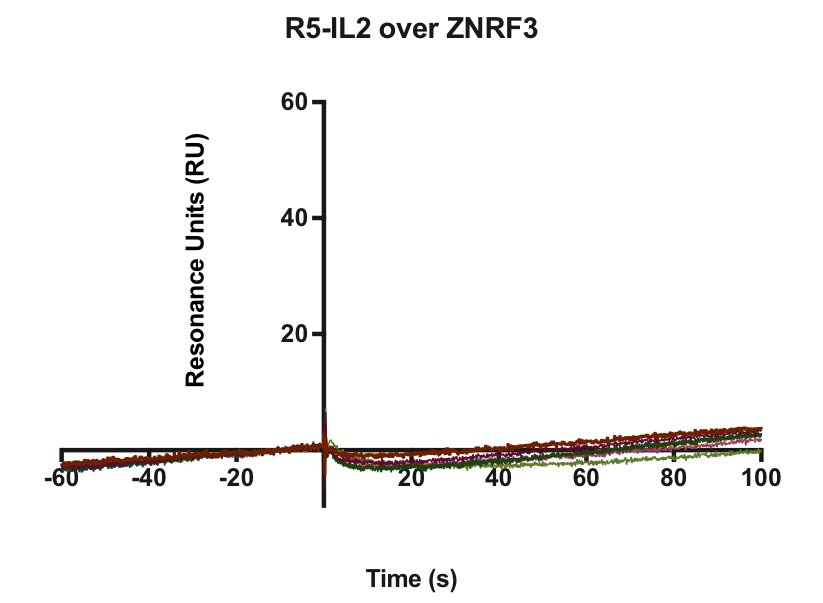

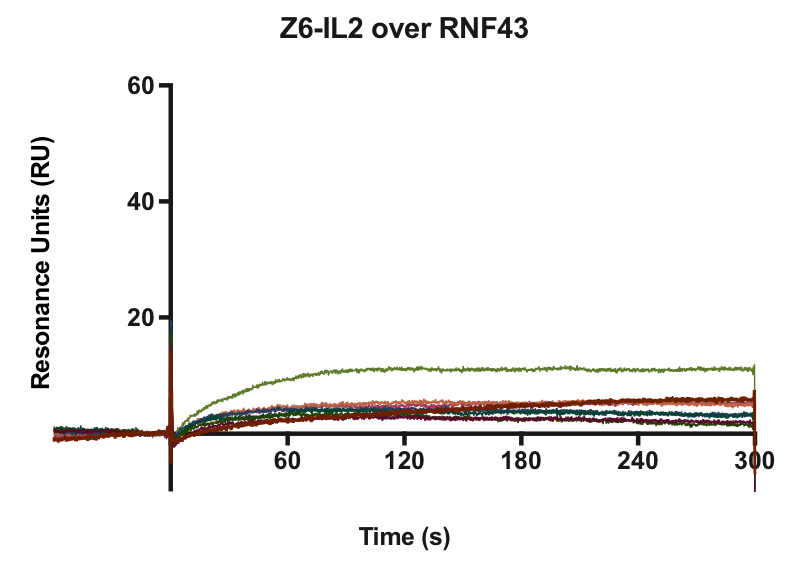

**Figure S3. Cross-reactivity of R5 and Z6 with RNF43 and ZNRF3.** SPR was used to assess whether the R5 scFv cross-reacts with ZNRF3, and whether the Z6 scFv cross-reacts with RNF43. Increasing concentrations of R5-IL2 (3-fold dilutions, green curve is maximum concentration of 1 μM) were injected over a surface coated with the ZNRF3 ECD (left) and increasing concentrations of Z6-IL2 (2-fold dilutions, green curve is maximum concentration of 1 μM) were injected over a surface coated with the RNF43 ECD (right). Injections were performed at T = 0 seconds. R5-IL2 was flowed over the surface for 100 seconds and Z6-IL2 was flowed over the surface for 300 seconds. In both cases no substantial binding was observed, indicating that neither scFv is cross-reactive.

S-4

Kd: 5.7nM

Kd: 8.4nM

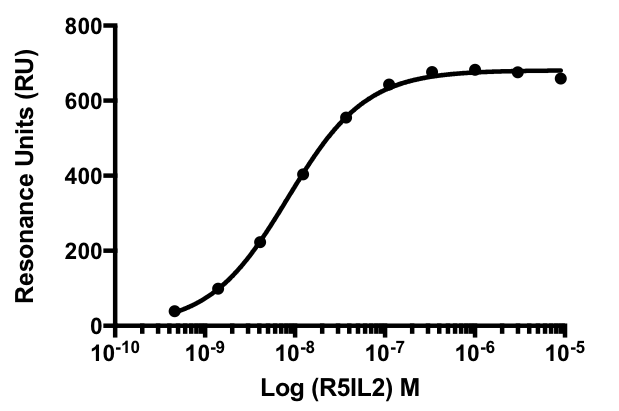

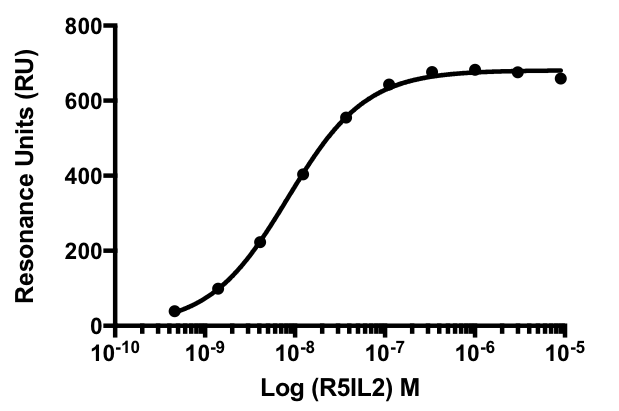

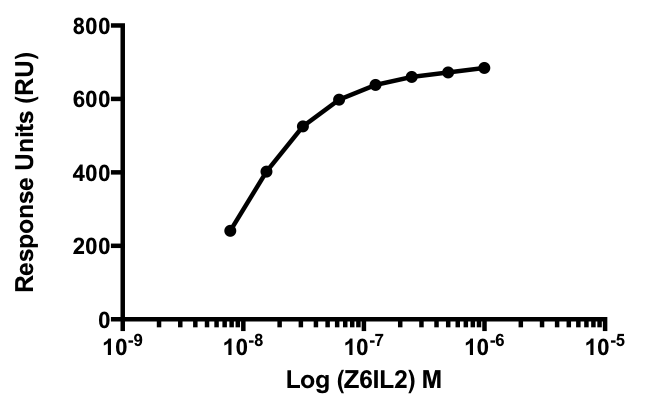

**Figure S4. Binding affinity between surrogate RSPOs and CD25.** SPR was used to determine the binding affinity between R5-IL2 or Z6-IL2 and CD25. Increasing concentrations of R5-IL2 (left) or Z6-IL2 (right) were injected over a surface coated with the ECD of CD25. The maximal RU values for each curve were plotted and the binding isotherms were fitted to a 1:1 model to determine the K_d_ values indicated on the plots.

S-5
